# BrainSpace: a toolbox for the analysis of macroscale gradients in neuroimaging and connectomics datasets

**DOI:** 10.1101/761460

**Authors:** Reinder Vos de Wael, Oualid Benkarim, Casey Paquola, Sara Lariviere, Jessica Royer, Shahin Tavakol, Ting Xu, Seok-Jun Hong, Sofie L. Valk, Bratislav Misic, Michael P. Milham, Daniel S. Margulies, Jonathan Smallwood, Boris C. Bernhardt

**Author notes:** Corresponding author Email address (Boris C. Bernhardt). Authors contributed equally to this work (Reinder Vos de Wael, Oualid Benkarim, Jonathan Smallwood, Boris C. Bernhardt).

## Abstract

Understanding how higher order cognitive function emerges from the underlying brain structure depends on quantifying how the behaviour of discrete regions are integrated within the broader cortical landscape. Recent work has established that this macroscale brain organization and function can be quantified in a compact manner through the use of multivariate machine learning approaches that identify manifolds often described as cortical gradients. By quantifying topographic principles of macroscale organization, cortical gradients lend an analytical framework to study structural and functional brain organization across species, throughout development and aging, and its perturbations in disease. More generally, its macroscale perspective on brain organization offers novel possibilities to investigate the complex relationships between brain structure, function, and cognition in a quantified manner. Here, we present a compact workflow and open-access toolbox that allows for (i) the identification of gradients (from structural or functional imaging data), (ii) their alignment (across subjects or modalities), and (iii) their visualization (in embedding or cortical space). Our toolbox also allows for controlled association studies between gradients with other brain-level features, adjusted with respect to several null models that account for spatial autocorrelation. The toolbox is implemented in both Python and Matlab, programming languages widely used by the neuroimaging and network neuroscience communities. Several use-case examples and validation experiments demonstrate the usage and consistency of our tools for the analysis of functional and microstructural gradients across different spatial scales.

## 1. Introduction

Over the last century, neuroanatomical studies in humans and non-human animals have highlighted two complementary features of neural organization. On the one hand, studies have demarcated structurally homogeneous areas with specific connectivity profiles, and ultimately distinct functional roles (e.g., Brodmann, 1909; von Economo and Koskinas, 1925; Palomero-Gallagher and Zilles, 2017; Flechsig, 1920; Glasser et al., 2016). In parallel, neuroanatomists have established that cortical organization may also be characterized in the form of smooth transitions at the system level, for example in terms of their histological properties and connectivity patterns (Von Bonin and Bailey, 1947; Bailey and Von Bonin, 1951; Goulas et al., 2019; Bailey and Von Bonin, 1951; Pandya et al., 2015). In a seminal attempt to synthesize an integrated view of the functions of the mammalian cortical landscape, Marsel Mesulam postulated a hierarchical sensory-fugal axis of cortical microstructural organization and connectivity (Mesulam, 1998). In this model, neural function is hypothesised to emerge not simply from modular neural systems, but through the complex interactions between regions. For example, peripheral neural systems such as primary sensory and motor regions, support functional interactions with the external world in a reasonably direct manner, while transmodal cortices, that take as input signals from regions towards the periphery, support increasingly abstract, and perceptually decoupled, cognitive operations (Mesulam, 1998).

Although much of the more recent work linking measures of neural processing (for example from functional magnetic resonance imaging, MRI) to cognition has focused on identifying discrete regions and modules and their specific functional roles (Eickhoff et al., 2018), recent conceptual and methodological developments have provided both the data and methods that allow macroscale brain features mapped to low dimensional manifold representations, also described as gradients (Margulies et al., 2016). Gradient analyses operating on connectivity data were applied to diffusion MRI tractography data in spefific brain regions (Cerliani et al., 2012; Bajada et al., 2017) as well as resting-state functional MRI connectivity maps (Margulies et al., 2016; Vos de Wael et al., 2018; Larivière et al., 2019; Tian and Zalesky, 2018; Marquand et al., 2017; Haak et al., 2018; Przeździk et al., 2019). Similar techniques have also been used to describe myelin-sensitive tissue measures and other morphological characteristics (Huntenburg et al., 2017; Paquola et al., 2019; Wagstyl et al., 2015). Other studies have used a similar approach to describe task based neural patterns either using meta-analytical co-activation mapping (Vos de Wael et al., 2018) or large-scale functional MRI task data sets (Shine et al., 2019). Gradients have been also successfully derived from non-imaging data that were registered to stereotaxic space, including hippocampal post mortem gene expression information (Vogel et al., 2019) and 3D histology data (Paquola et al., 2019), to explore cellular and molecular signatures of neuroimaging and connectome measures. Core to these techniques is the computation of an affinity matrix that captures inter-area similarity of a given feature followed by the application of dimensionality reduction techniques to identify a gradual ordering of the input matrix in a lower dimensional manifold space (Figure 1).

**Figure 1:**
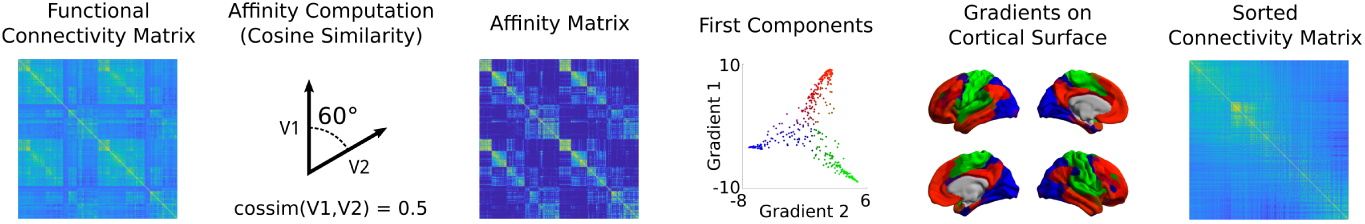
A typical gradient identification workflow. Starting from an input matrix (here, functional connectivity), we use a kernel function to build the affinity matrix (here capturing the connectivity of each seed region). This matrix is decomposed, often via linear rotations or non-linear manifold learning techniques into a set of principal eigenvectors describing axes of largest variance. The scores of each seed onto the first two axes are shown in the scatter plot, with colors denoting position in this 2D space. These colors may be projected back to the cortical surface and the scores can be used to sort the input connectome.

The ability to describe the brain wide neural activity in a single manifold offers the possibility to understand how the integrated nature of neural processing gives rise to function and dysfunction. Adopting a macroscale perspective on cortical organization has already provided important insights into how cortex-wide patterns relate to cortical dynamics (Wang et al., 2019) and high level cognition (Sormaz et al., 2018; Murphy et al., 2018, 2019; Shine et al., 2019). Furthermore, several studies have leveraged gradients as an analytical framework to describe atypical macroscale organization across clinical conditions, for example, by showing perturbations in functional connectome gradients in autism (Hong et al., 2019) and schizophrenia (Tian et al., 2019). Finally, comparisons of gradients across different imaging modalities have highlighted the extent to which structure directly constrains functional measures (Paquola et al., 2019), while consideration of gradients across species has highlighted how evolution has shaped more integrative features of the cortical landscape (Goulas et al., 2019; Buckner and Krienen, 2013; Huntenburg et al., 2018; Xu et al., 2019; Buckner and Margulies, 2019; Fulcher et al., 2019).

The growth in our capacity to map whole brain cortical gradients, coupled with the promise of a better understanding of how structure gives rise to function, highlights the need for a set of tools that support the analysis of neural manifolds in a compact and reproducible manner. The goal of this paper is to present an open-access set of easy-to-use tools that allow the identification, visualization, and analysis of macroscale gradients of brain organization. We hope this will provide a method for calculating cortical manifolds that facilitates their use in future empirical work, allows comparison between studies, and allows for result replicability. To offer flexibility in implementation, we provide our toolbox in both Python and Matlab, two languages widely used in the neuroimaging and network neuroscience communities. Associated functions are freely available for download and complemented with an expandable online documentation. We anticipate that our toolbox will help researchers interested in studying gradients of cortical organization, and propel further work that establishes the overarching principles through which structural and functional organization of human and non-human brains gives rise to key aspects of human cognition.

## 2. Methodology

### 2.1. Data and code availability statement

Our toolbox is openly available at: http://github.com/MICA-MNI/BrainSpace. The toolbox contains a parallel python and matlab implementation with closely-matched functionality. Along with the code, the toolbox contains several surface models, parcellations across multiple scales, and example data to reproduce the evaluations presented in this tutorial. Additionally, documentation of the pro-posed toolbox is available for both python and matlab implementations via http://brainspace.readthedocs.io.

### 2.2. Input data description

Our toolbox requires a real input matrix. Let *X* ∈ ℝ^*n*×*p*^ be a matrix aggregating features of several seed regions. In other words, each seed is represented by a *p*-dimensional vector, **x**_*i*_, built based on the features of the *i*-th seed region, where 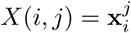 denotes the *j*-th feature of the *i*-th seed. In many neuroimaging applications, *X* may represent a connectivity metric (e.g., resting-state functional MRI connectivity or diffusion MRI tractography derived structural connectivity) between different seed and target brain regions. When seed and target regions are identical, the input matrix *X* is square. Furthermore, if the connectivity measure used to build the matrix is non-directional, *X* is also symmetric. If seeds and targets are different, for example when assessing connectivity patterns of a given region with the rest of the brain (Vos de Wael et al., 2018; Haak et al., 2018), we may have that *n* ≠ *p*, resulting in a non-square matrix. The dimensions and symmetry properties of the input matrix *X* may interact with the dimensionality reduction procedures presented in section 2.4. A simple strategy to make matrices symmetric and squared is to use kernel functions, which will be covered in the following section.

### 2.3. Affinities and kernel functions

Since we are interested in studying the relationships between the seed regions in terms of their features (e.g., connectivity with target regions), our toolbox provides several kernel functions to compute the relationship between every pair of seed regions and derive a non-negative square symmetric affinity matrix *A* ∈ ℝ^*n*×*n*^, where *A*(*i, j*) = *A*(*j, i*) denotes the similarity or affinity between seeds *i* and *j*. Moreover, a square symmetric matrix is a requirement for the next step in our framework (i.e., dimensionality reduction). Note that when the input matrix *X* is already square and symmetric (e.g., seed and target regions are the same), there may be no need to derive the affinity matrix and *X* can be used directly to perform dimensionality reduction. Accordingly, BrainSpace provides the option of skipping this step and using the input matrix as the affinity.

There are numerous approaches to build affinity matrices. Our toolbox currently implements the following kernels: Gaussian, cosine similarity, normalized angle similarity, Pearson’s correlation coefficient, and Spearman rank order correlations. For simplicity, let **x** = **x**_*i*_ and **y** = **x**_*j*_, these kernels can be expressed as follows:

1. Gaussian kernel:

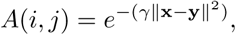

where *γ* is the inverse kernel width and ‖ ·‖ denotes the *l*_2_-norm.
2. Cosine similarity:

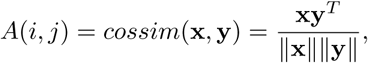

where *cossim*(·,·) is the cosine similarity function and *T* stands for transpose.
3. Normalized angle similarity:

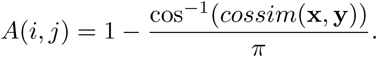
4. Pearson correlation:

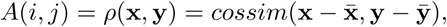

where *ρ* is the Pearson correlation coefficient, and 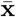 and 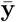 denote the means of **x** and **y**, respectively.
5. Spearman rank order correlation:

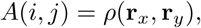

where **r**_*x*_ and **r**_*y*_ denote the ranks of **x** and **y**, respectively.

While not being exhaustive, the first version of BrainSpace thus includes commonly used kernels in the gradient literature and additional ones for experimentation. To our knowledge, no gradient paper has used Pearson or Spearman correlation. Note that if *X* is already row-wise demeaned, Pearson correlation amounts to cosine similarity. The Gaussian kernel is widely used in the machine learning community (for example in the context of Laplacian eigenmaps and support vector machines), which provides a simple approach to convert Euclidean distances between our seeds into similarities. Cosine similarity, [example application (Margulies et al., 2016)], computes the angle between our feature vectors to describe their similarity. It is important to note that cosine similarity ranges between −1 and 1, with negative correlations to be transformed to non-negative values. This motivated the inclusion of the normalized angle kernel, [example application: (Vos de Wael et al., 2018)], as it was developed to circumvent negative similarities by transforming the similarities to [0, 1], with 1 denoting identical angles, and 0 opposing angles. Still, cosine similarity and Pearson and Spearman correlation coefficients may produce negative values (i.e., [−1, 1]) if the feature vectors are negatively correlated. A simple approach to deal with this issue is to set negative values to zero. Additionally, BrainSpace provides an option for column-wise thresholding of the input matrix as done previously (Margulies et al., 2016; Vos de Wael et al., 2018; Paquola et al., 2019; Hong et al., 2019). Each vector of the input matrix is thresholded at a given sparsity (e.g., by keeping the weights of the top 10% entries for each region). This procedure ensures that only strong, and potentially less noisy, connections contribute to the gradient.

In addition to the aforementioned kernels, BrainSpace provides the option of skipping this step and using the input matrix as an affinity matrix as well as the option to provide a custom kernel.

### 2.4. Dimensionality reduction

Recall that in the input matrix, each seed **x**_*i*_ is defined by a *p*-dimensional feature vector, where *p* may denote hundreds of parcels or thousands of vertices/voxels. The aim of dimensionality reduction techniques is to find a meaningful underlying low-dimensional representation, 𝒢 ∈ ℝ^*n*×*m*^ with *m* ≪ *p*, hidden in the high-dimensional ambient space. These methods can be grouped into linear and non-linear techniques. The former use a linear transformation to unravel the latent representation, while techniques in the second category use non-linear transformations. BrainSpace provides three of the most widely used dimensionality reduction techniques for macroscale gradient mapping: principal component analysis (PCA) for linear embedding, and laplacian eigenmaps (LE) and diffusion mapping (DM) for non-linear dimensionality reduction.

1. PCA is a linear approach that transforms the data to a low-dimensional space represented by a set of orthogonal components that explain maximal data variance. Given a column-wise demeaned version of the input matrix *X*_*d*_, the low-dimensional representation is computed as follows:

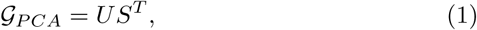

where *U* are the left singular vectors and *S* a diagonal matrix of singular values obtained after factorizing the input matrix using singular value decomposition (SVD), *X*_*d*_ = *USV*^*T*^. Note that, although here we present the SVD version, PCA can also be performed based on the eigendecomposition of the covariance matrix of *X*, for example.
2. LE is a non-linear dimensionality reduction technique that uses the graph Laplacian of the affinity matrix *A* to perform the embedding:

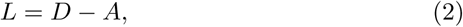

where the degree matrix *D* is a diagonal matrix defined as *D*(*i, i*) = Σ_*j*_ *A*(*i, j*) and *L* is the graph Laplacian matrix. Note that we can also work with its normalized version instead *L*_*s*_ = *D*^−1^*/*^2^*LD*^−1^*/*^2^ (Ng et al., 2001). LE then proceeds to solve the generalized eigenvalue problem:

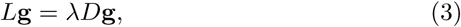

where the eigenvectors **g**_*k*_ corresponding to the *m* smallest eigenvalues *λ*_*k*_ (excluding the first eigenvalue) are used to build the new low-dimensional representation:

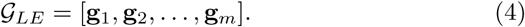
3. DM also seeks a non-linear mapping of the data based on the diffusion operator *P*_*α*_, which is defined as follows:

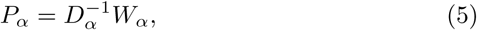

where *α* ∈ [0, 1] is the anisotropic diffusion parameter used by the diffusion operator, *W*_*α*_ = *D*^−1*/α*^*AD*^−1*/α*^ is built by normalizing the affinity matrix according to the diffusion parameter and *D*_*α*_ is the degree matrix derived from *W*_*α*_. When *α* = 0, the diffusion amounts to normalized graph Laplacian on isotropic weights, for *α* = 1, it approximates the Laplace-Beltrami operator and for the case where *α* = 0.5 it approximates the Fokker-Planck diffusion (Coifman and Lafon, 2006). It controls the influence of the density of sampling points on the manifold (*α* = 0, maximal influence; *α* = 1, no influence). In the gradient literature, the anisotropic diffusion hyper-parameter is commonly set to *α* = 0.5 (Margulies et al., 2016; Vos de Wael et al., 2018; Larivière et al., 2019), a choice that retains the global relations between data points in the embedded space. Similarly to LE, DM computes the eigenvalues and eigenvectors of the diffusion operator. However, in this case, the new representation is constructed with the scaled eigenvectors corresponding to the largest eigen-values, after omitting the eigenvector with the largest eigenvalue:

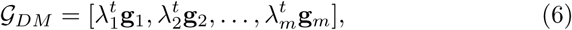

where *t* is the time parameter that represent the scale.

### 2.5. Alignment of gradients

Gradients computed separately for two or more datasets (e.g., patients vs controls, left vs right hippocampi) may not be directly comparable due to different eigenvector orderings in case of eigenvalue multiplicity (i.e., eigenvalues with the same value) and sign ambiguity of the eigenvectors (Lombaert et al., 2013). Aligning gradients improves comparability and correspondence. However, we recommend always visually inspecting the output of gradient alignments; if the manifold spaces are substantially different then alignments may not provide sensible output. In the first version of the BrainSpace toolbox, gradients can be aligned using Procrustes analysis (Langs et al., 2015) or implicitly by joint embedding.

#### Procrustes analysis

Briefly, given a source 𝒢_*s*_ and a target 𝒢_*t*_ representations, Procrustes analysis seeks an orthogonal linear transformation *ψ* to align the source representation to the target, such that *ψ*(𝒢_*s*_) and 𝒢_*t*_ are superimposed. Translation and scaling can also be performed by initially centering and normalizing the data prior to finding the transformation. For multiple datasets, a generalized Procrustes analysis is employed. Let 𝒢_*k*_, *k* = 1, 2,…, *N* be the low-dimensional representations of *N* different datasets (i.e., input matrices *X*_*k*_). The procedure proceeds iteratively by aligning all representations *G*_*k*_ to a reference and updating the reference 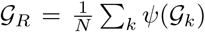 by averaging the aligned representations. In the first iteration, the reference can be chosen from the available representations, or an out-of-sample template can be provided (e.g., from a hold-out group).

#### Joint embedding

Joint embedding is a dimensionality reduction technique that finds a common underlying representation of multiple datasets by using a simultaneous embedding (Xu et al., 2019). The main challenge of this technique is to find a meaningful approach to establish correspondences between the original datasets (i.e., *X*). In the current version of BrainSpace, joint alignment is implemented based on spectral embedding and it is available for LE and DM. The only difference with these methods is that the embedding, rather than using the affinity matrices individually, is based on the joint affinity matrix 𝒥, which is built as:

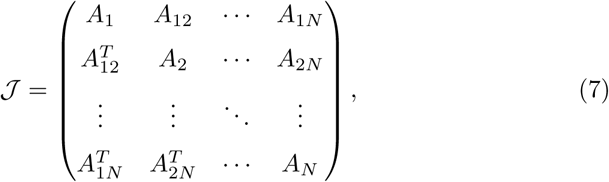

where *A*_*k*_ is the intra-dataset affinity of input matrix *X*_*k*_ and *A*_*ij*_ is the interdataset affinity between *X*_*i*_ and *X*_*j*_. As of the current version, both sets of affinities are built using the same kernel. It is important to note, therefore, that joint embedding can only be used if the input matrices have the same features (e.g., identical target regions). After the embedding, the resulting shared representation 𝒢_*J*_ = [𝒢_1_, 𝒢_2_,…, 𝒢_*N*_]^*T*^ will be composed of *N* individual low-dimensional representations, such that for the *k*-th input matrix 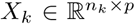, the corresponding representation is 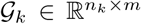, where *n*_*k*_ is the number of seeds (i.e., rows) of *X*_*k*_.

### 2.6. Null models

Many researchers may use gradients to compare axes of the brain to other continuous brain markers such as cortical thickness measures or estimates of myelination. Given the spatial autocorrelation present in many modalities, a linear regression or similar methods may provide biased test statistics. To circumvent this issue, we recommend comparing the observed test statistic to those of a set of distributions with similar spatial autocorrelation. To this end we provide two methods: spin permutations (Alexander-Bloch et al., 2018) and Moran spectral randomization (MSR) (Wagner and Dray, 2015; Dray, 2011). In cases where the input data lies on a surface and most of the sphere is used, we recommend using spin permutation otherwise we recommend MSR. When performing a statistical test with multiple gradients as either predictor or response variable, we recommend randomizing the non-gradient variable as these randomizations need not maintain statistical independence across different eigenvectors.

#### Spin permutations

Spin permutation analysis leverages the spherical representations of the cerebral cortex, such as those derived from FreeSurfer (Fischl, 2012) or CIVET (Kim et al., 2005), to address the problem of spatial autocorrelation in statistical inference. In short, spin permutations estimate the null distribution by randomly rotating the spherical projections of the cortical surface while preserving the spatial relationships within the data (Alexander-Bloch et al., 2018). Let *V* ∈ ℝ^*l*×3^ be the matrix of vertex coordinates in the sphere, where *l* is the number of vertices, and *R* ∈ ℝ^3×3^ a matrix representing a rotation along the three axes uniformly sampled from all possible rotations (Blaser and Fryzlewicz, 2016). The rotated sphere *V*_*r*_ is computed as follows:

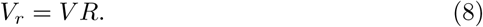

Samples of the null distribution are then created by assigning each vertex on *V*_*r*_ the data of its nearest neighbor on *V*.

#### Moran spectral randomization

Borrowed from the ecology literature, MSR can also be used to generate random variables with identical or similar spatial autocorrelation (in terms of Moran’s *I* correlation coefficient (Cliff and Ord, 1973)). This approach requires building a spatial weight matrix defining the relationships between the different locations. For our particular case, given a surface mesh with *l* vertices (i.e., locations), its topological information is used to build the spatial weight matrix *L* ∈ ℝ^*l*×*l*^, such that *L*(*i, j*) *>* 0 if vertices *i* and *j* are neighbors, and *L*(*i, j*) = 0 otherwise. In BrainSpace, *L* is built using the inverse distance between each vertex and the vertices in its immediate neighborhood, although other neighborhoods and weighting schemes (e.g., binary or Gaussian weights) could be used. Then, *L* is doubly centered and eigendecomposed into its whole spectrum, with the resulting eigenvectors ℳ ∈ ℝ^*l*×(*l*−1)^ being the so-called Moran eigenvector maps. Note that the eigenvector with 0 eigenvalue is dropped. The advantage of MSR is that we can work with the original cortical surfaces, and thus skip the potential distortions introduced by the spherical mesh parameterization.

In order to generate the null distributions, let **u** ∈ ℝ^*l*^ be an input feature vector defined on each vertex of our surface (e.g., cortical thickness), and **r** ∈ ℝ^*l*−1^ the correlation coefficients of **u** with each spatial eigenvector in ℳ. MSR aims to find a randomized feature vector **z** that respects the autocorrelation observed in **u** as follows:

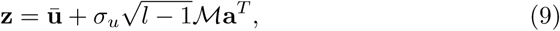

where **ū** and *σ*_*u*_ stand for mean and standard deviation of **u** respectively, and **a** ∈ ℝ^*l*−1^ is a vector of random coefficients. Three different methods exist for generating vector **a**: singleton, pair, and triplet procedures. Currently, only the singleton and pair procedures are supported in BrainSpace. Let **v** = ℳ**r**^*T*^, in the singleton procedure, **a** is computed by randomizing the sign of each element in **v** (i.e., **a**_*i*_ = ±**v**_*i*_). In the pair procedure, the elements of **v** are randomly changed in pairs. Let (**v**_*i*_, **v**_*j*_) be a pair of elements randomly chosen, then **a** is updated such that **a**_*i*_ = *q*_*ij*_ cos(*ϕ*) and **a**_*j*_ = *q*_*ij*_ sin(*ϕ*), with 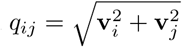 and *ϕ* ∼ 𝒰 (0, 2*π*) randomly drawn from a uniform distribution. If the number of elements of **v** is odd, the singleton procedure is used for the remaining element.

As opposed to the singleton procedure, the null data generated by the pair procedure does not fully preserve the observed spatial autocorrelation. We therefore recommend the singleton procedure, unless the number of required randomizations exceeds 2^*l*−1^, which is the maximum number of unique randomizations that can be produced using the singleton procedure.

## 3. Examples and evaluations

In this section, we demonstrate the usage of our toolbox through several examples on gradient mapping and null model generation. Evaluations are based on a subset of the Human Connectome Project (HCP) dataset (Van Essen et al., 2013), as in prior work (Vos de Wael et al., 2018). Matlab code is presented in the main version of the paper. The python version is available in Appendix A.

### 3.1. Generating gradients

To first illustrate the basic functionality of the toolbox, we computed gradients derived from resting-state functional MRI functional connectivity (FC). In short, the input matrix was made sparse (to 10% sparsity) and a cosine similarity matrix was computed. Next, we applied the three different manifold algorithms (i.e., PCA, LE, DM) and plot their first and second gradients on the cortical surface (Sample Code 1). Resulting gradients (Figure 2) derived from all dimensionality reduction techniques resemble those published previously (Margulies et al., 2016), although for PCA the somatomotor to visual gradient explains more variance than the default mode to sensory gradient.

**Figure 2:**
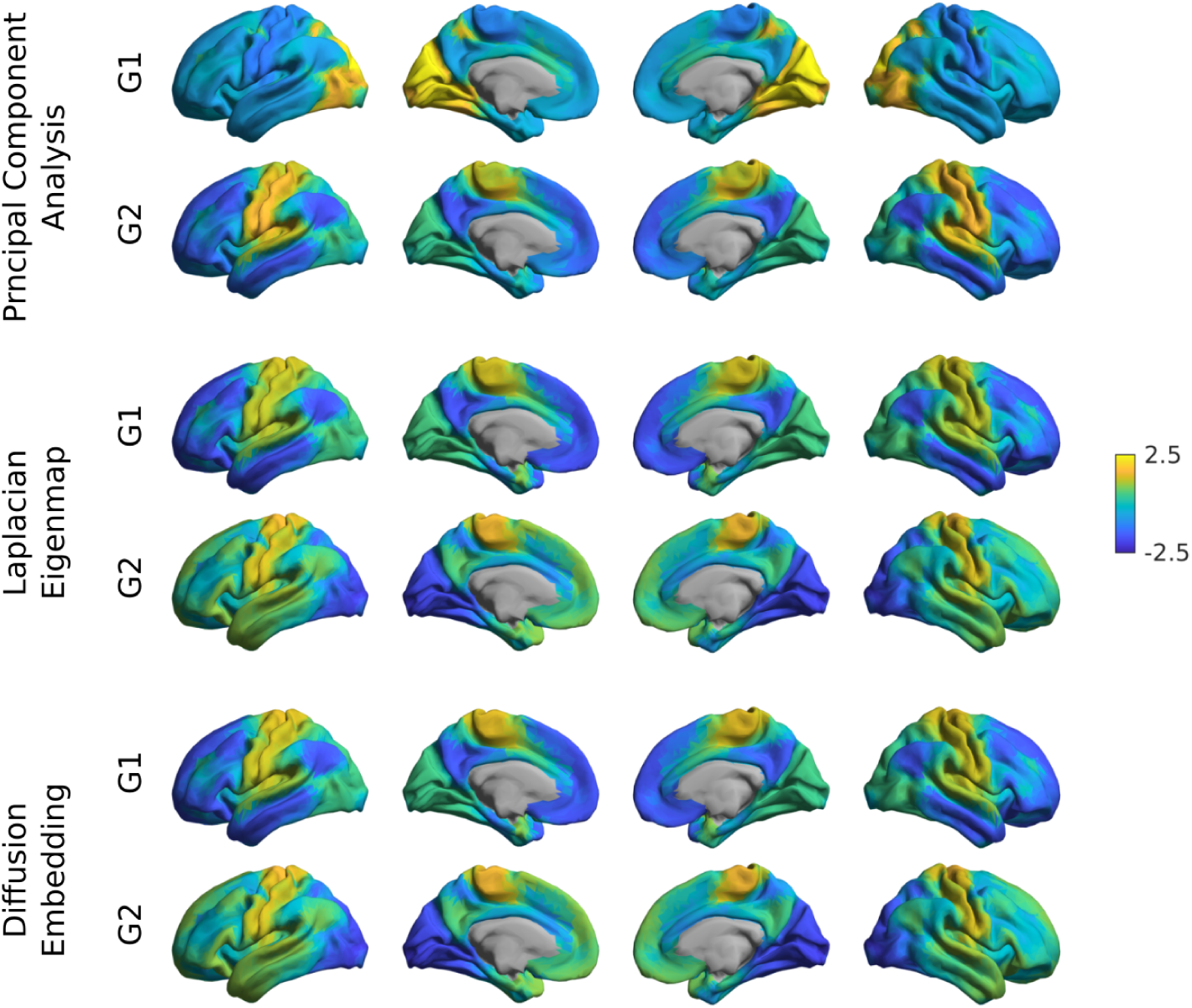
Gradient construction with different dimensionality reduction techniques. Gradient 1 (G1) and 2 (G2) of FC were computed using a cosine similarity affinity computation, followed by either PCA, LE, or DM. Gradients were z-scored before plotting.

Sample Code 1: A minimal Matlab example for plotting the first gradient of an input data matrix on the cortical surface. Equivalent Python code is provided in sample code 5 in Appendix A.

~~~
*% Create a GradientMaps object*
G = GradientMaps(‘kernel’,’cosine’, ‘approach’,’dm’);
*% Apply GradientMaps to the data*
G = G.fit(data_matrix);
*% Load surfaces*
left_surface = read_surface(‘left_surface_file.obj’);
right_surface = read_surface(‘right_surface_file.obj’);
*% Plot first gradient on the cortical surface*
plot_hemispheres(G.gradients{1}(:,1), {left_surface,right_surface})
~~~

### 3.2. Aligning gradients

#### a) Gradient alignment across modalities

Based on subjects present in both the FC dataset as well as those used in the validation group of (Paquola et al., 2019), we assessed the correspondence between gradients computed from different modalities and evaluated increases in correspondence when gradient alignment was used. The two modalities that were evaluated are FC and microstructural profile covariance (MPC). Here we compare gradients of the two modalities in an unaligned form, as well as after Procrustes alignment and joint embedding (Sample Code 2, Figure 3). Gradient correspondence increases following Procrustes alignment and a marked increase following joint embedding. Beyond maximizing correspondence, the use of Procrustes vs joint embedding can also depend on the specific applications. Procrustes alignment preserves the overall shape of the different gradients and can thus be a preferable approach to compare different gradients. Joint embedding, on the other hand, identifies a joint solution that maximizes their similarity, so the resulting gradient may be more of a ‘hybrid’ of the input manifolds. Joint embedding is thus a technique to identify correspondence and map from one space to another, and is conceptually related to widely used multivariate associative techniques such as canonical correlation analysis or partial least squares which seek to maximize the linear associations between two multidimensional datasets (Mcintosh and Misic, 2013). Note that the computational cost of joint embedding is substantially higher, so Procrustes analysis may be the preferred option when computational resources are a limiting factor.

**Figure 3:**
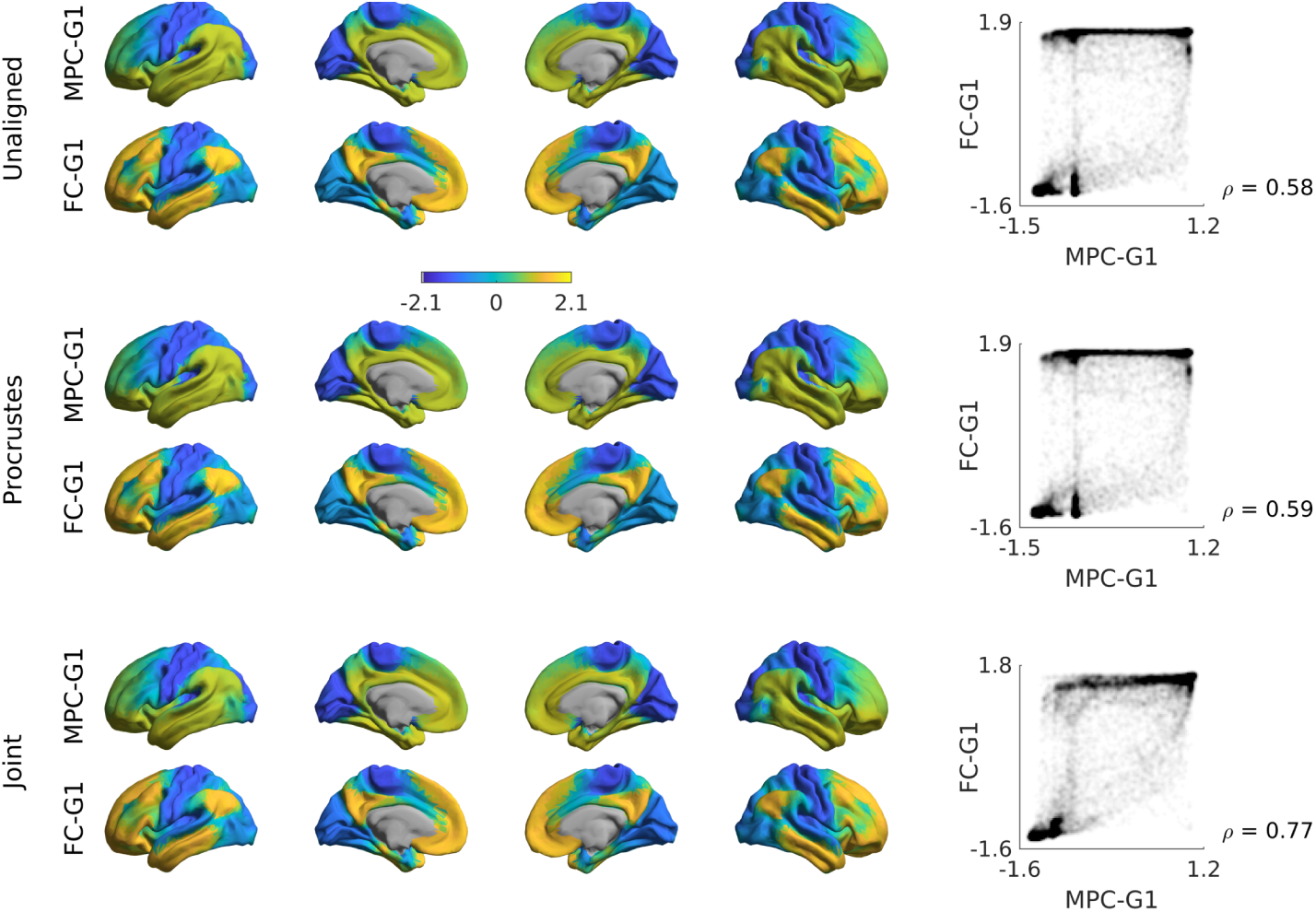
Comparison of alignment methods across modalities. Unaligned gradients 1 (top) of MPC and FC were derived using cosine similarity and diffusion embedding. Alignments using Procrustes analyses (middle) and joint embedding (bottom) are also shown. Smoothed scatter plots show a moderate increase in Spearman correlation after joint embedding. Gradients were z-scored before plotting.

Sample Code 2: A minimal Matlab example for creating and plotting gradients from different modalities, with different alignments. Equivalent Python code is provided in sample code 6 in Appendix A.

~~~
*% Create two GradientMaps objects with different alignments*
Gp = GradientMaps(‘kernel’,’cosine’, ‘approach’,’dm’, *…*
‘alignment’,’procrustes’);
Gj = GradientMaps(‘kernel’,’cosine’, ‘approach’,’dm’, *…*
‘alignment’,’joint’);
*% Apply GradientMaps to the data*
Gp = Gp.fit({mpc,fc});
Gj = Gj.fit({mpc,fc}});
*% Load surfaces*
left_surface = read_surface(‘left_surface_file.obj’);
right_surface = read_surface(‘right_surface_file.obj’);
*% Plot first MPC gradient of Procrustes alignment*
plot_hemispheres(Gp.aligned{1}(:,1), {left_surface,right_surface});
*% Plot first FC gradient of joint embedding*
plot_hemispheres(Gj.aligned{2}(:,1), {left_surface,right_surface});
~~~

#### b) Gradient alignment across individuals

Researchers may also be interested in comparing gradient values between individuals (Hong et al., 2019), for example to assess perturbations in FC gradients as a measure of brain network hierarchy. One possible approach could be to first build a group-level gradient template, to which both diagnostic groups are aligned using Procrustes rotation. After that, the two groups can be compared statistically and at each vertex, for example using the SurfStat toolbox (Worsley et al., 2009) or alternative tools. In the example below, we present the steps to derive, align, and compare the principal functional gradient between a cohort of individuals. We computed a template gradient from an out-of-sample dataset of 134 subjects from the HCP dataset (the validation cohort used by (Vos de Wael et al., 2018)). Next, we used Procrustes analysis to align individual’s gradients of each subject to the group level template (Sample Code 3, Figure 4).

**Figure 4:**
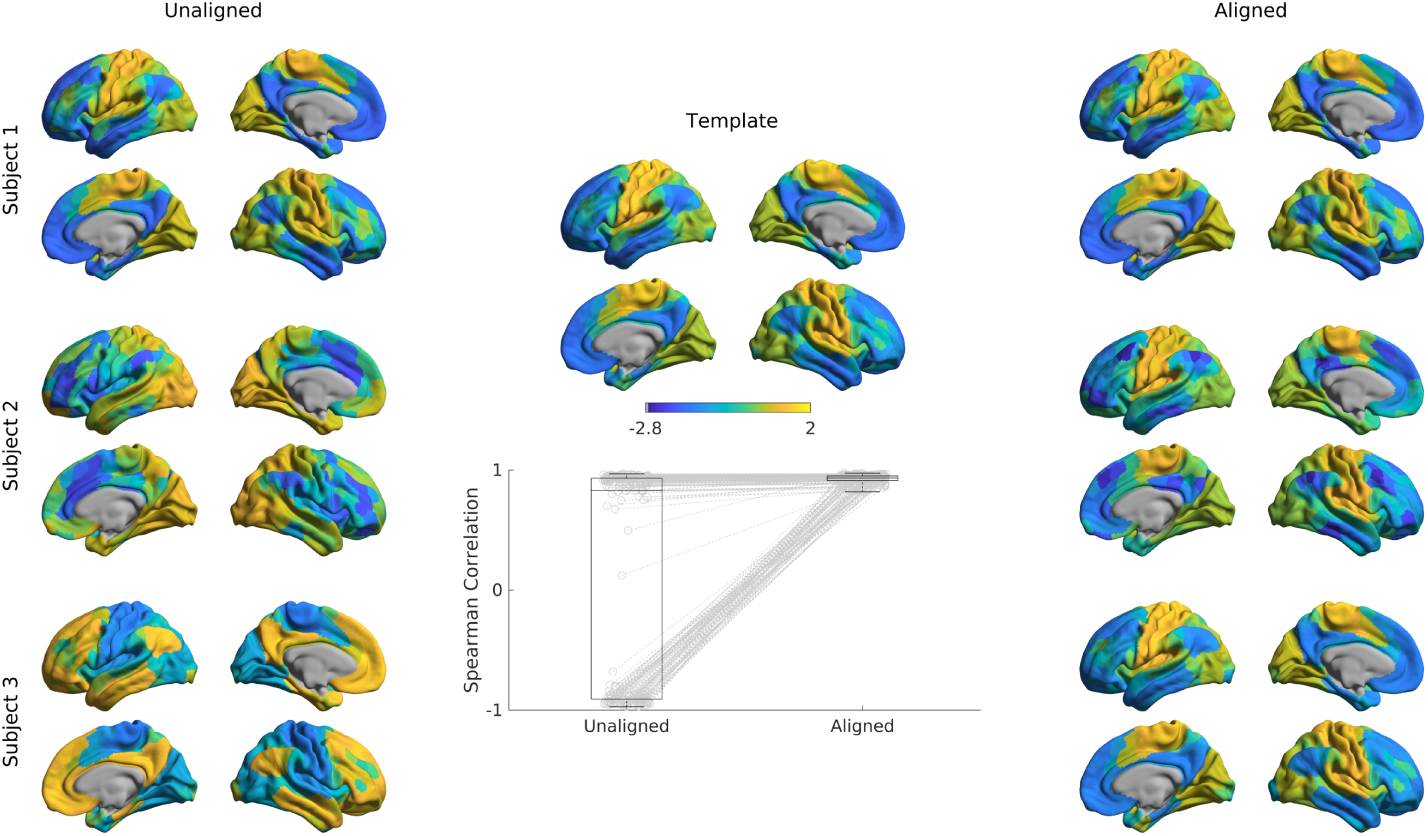
Alignment of subjects to a template. Gradients 1 of single subjects computed with the cosine similarity kernel and diffusion embedding manifold (left) were aligned to an out-of-sample template (middle) using Procrustes Analysis, creating the aligned gradients (right). Box-plot shows the Spearman correlations of each subjects’ gradient 1 to the template gradient 1 both before, and after Procrustes alignment

Sample Code 3: A minimal Matlab example for aligning the gradients of two individuals to a template gradient. Equivalent Python code is provided in sample code 7 in Appendix A.

~~~
*% Create a GradientMaps object for the template*
Gt = GradientMaps(‘kernel’,’cosine’, ‘approach’,’dm’);
*% Apply GradientMaps to template data*
Gt = Gt.fit(template_data);
*% Create a GradientMaps object for the individuals*
Gs = GradientMaps(‘kernel’,’cosine’, ‘approach’,’dm’, *…*
‘alignment’,’procrustes’);
*% Compute gradients of all subjects and align to template*
Gs = Gs.fit({subject1_data,subject2_data}, *…*
‘reference’,Gt.gradients{1});
*% Load surfaces*
left_surface = read_surface(‘left_surface_file.obj’);
right_surface = read_surface(‘right_surface_file.obj’);
*% Plot the first aligned gradient for subject 2*
plot_hemispheres(Gs.aligned{2}(:,1), {left_surface,right_surface});
~~~

### 3.3. Gradients across different spatial scales

The gradients presented so far were all derived at a vertex-wise level, which requires considerable computational resources. To minimize time and space requirements and to make results comparable to other parcellation-based studies, some users may be interested in deriving gradients using parcellated data. To illustrate the effect of using different parcellations, we repeated the gradient identification analysis across different spatial scales for both a structural and a functional parcellation. Specifically, we subdivided the conte69 surface into 400, 300, 200, and 100 parcels based on both a clustering of a well-established anatomical atlas (Desikan et al., 2006), as well as a recently published local-global functional clustering (Schaefer et al., 2017) and built FC gradients from these representations (Figure 5). These parcellations and subsampling schemes are provided in the *shared* folder of the BrainSpace toolbox.

**Figure 5:**
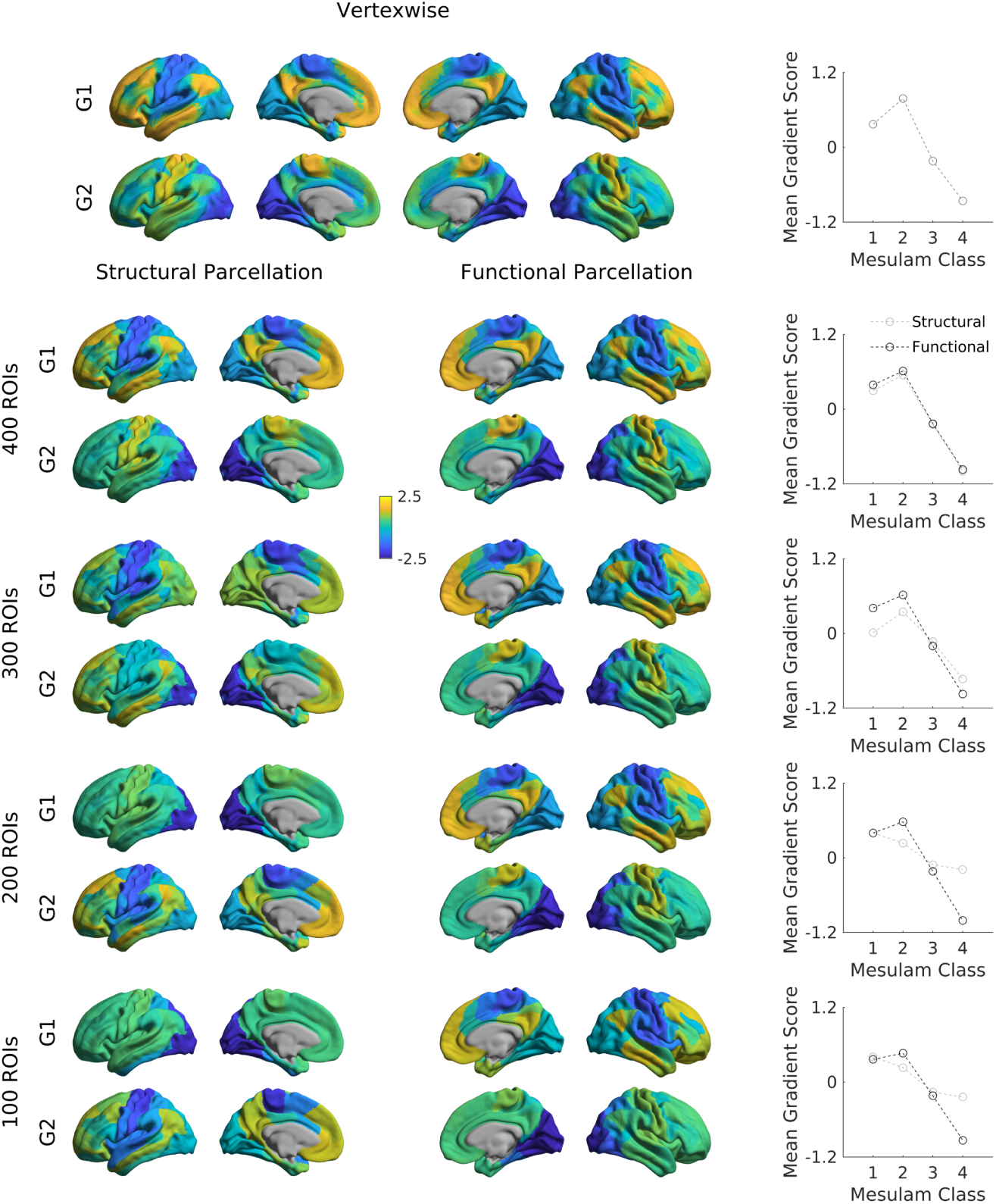
Functional gradients across spatial scales. The cortex was subdivided into 100 (first row), 200 (second row), 300 (third row), and 400 (fourth row) regions of interest based on an anatomical (left) and functional (right) parcellation. Displayed are gradients 1 (G1), and 2 (G2), each for one hemisphere only. Line plots show the average gradient score within each Mesulam class for the functional (dark gray) and structural (light gray).

Overall, with increasing spatial resolution the FC gradients become more pronounced and gradients derived from functional and structural parcellations were more similar. At a scale of 200 nodes or lower, putative functional boundaries may not be as reliably captured when using anatomically-informed parcellations, resulting in more marked alterations in the overall shape of the gradients.

For further evaluation, we related the above gradients across multiple scales relative to Mesulam’s classic scheme of laminar differentiation and hierarchy (Paquola et al., 2019). It shows clear correspondence between the first gradient and the Mesulam hierarchy for high resolution data of 300 nodes and more, regardless of the parcellation scheme. While high correspondence was still seen for functional parcellations at lower granularity, it was markedly reduced for structural. For researchers interested in using Mesulam’s parcellation, it has been provided on Conte69 surfaces in the *shared* folder of the BrainSpace toolbox. We also provide the evaluated FC and MPC gradients across these different spatial scales. Such gradients can be used to, for example stratify other imaging measures, including functional activation and connectivity patters (Hong et al., 2019; Murphy et al., 2018), meta-analytical syntheses (Murphy et al., 2019; Margulies et al., 2016), cortical thickness measures or Amyloid-beta PET uptake data (Lowe et al., 2019).

### 3.4. Null models

Here we present an example to assess the significance of correlations between FC gradients and data from other modalities (cortical thickness and T1w/T2w image intensity in this example). We present code to generate previously proposed spin tests (Alexander-Bloch et al., 2018), which preserve the auto-correlation of the permuted feature(s) by rotating the feature data on the sphere. In our example (Figure 6), one can clearly see that correlations between FC gradients and T1w/T2w stay highly significant even when comparing the correlation to 1000 null models whereas correlations between FC gradients and cortical thickness appear non-significant.

**Figure 6:**
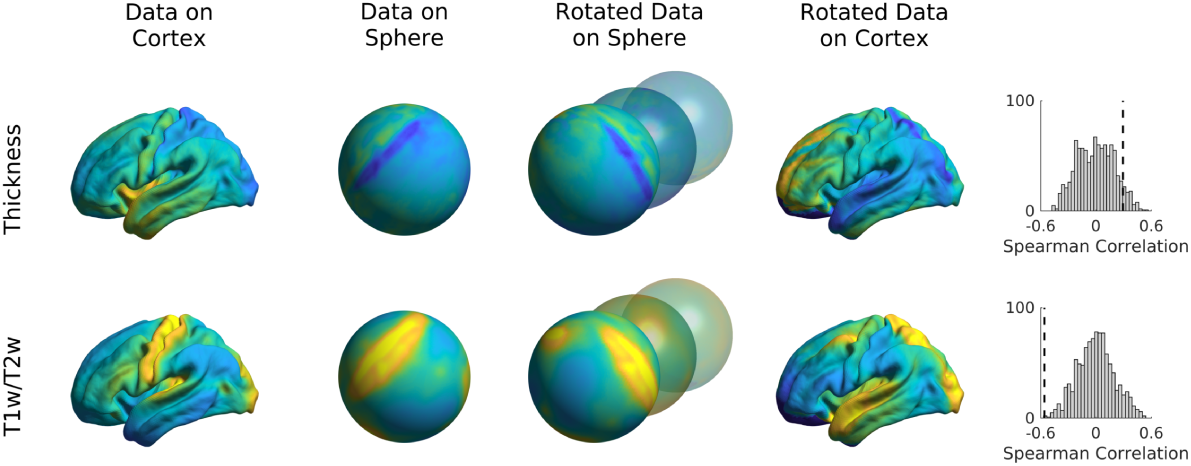
Spin tests of cortical thickness and t1w/t2w intensity. Data were rotated on the sphere 1000 times and the Spearman correlation between FC gradient 1 and the rotated data were computed. Distribution of correlations are shown in the histograms with the dashed lines denoting the true correlation.

Sample Code 4: A minimal Matlab example for building null models based on spin tests. Equivalent Python code is provided in sample code 8 in Appendix A.

~~~
*% Load spheres*
left_sphere = read_surface(‘left_sphere_file.obj’);
right_sphere = read_surface(‘right_sphere_file.obj’);
*% Generate 1000 spin permutations*
n_perm = 1000;
features_spin = spin_permutations({left_features,right_features}, *…*
{left_sphere,right_sphere},n_perm);
~~~

## 4. Discussion

While tools for unsupervised manifold identification and their alignment are widely available and extensively used in data science across multiple research domains^2^, and while some prior studies made their workflow openly accessible (see (Margulies et al., 2016; Haak et al., 2018; Guell et al., 2019) ^3 4 5^) we currently lack a unified software package that incorporates the major steps of gradient construction and evaluation for neuroimaging and connectome analysis datasets. We aimed to fill this gap with BrainSpace, a compact toolbox for the identification and analysis of low-dimensional gradients for any given regional or connectome-level feature. As such, the toolbox provides a simple entry point for researchers interested in studying gradients as windows into brain organization and function. Tools are available in Python and Matlab, two widely used programming languages in the neuroimaging and connectomics communities, and openly shared via Github at http://github.com/MICA-MNI/BrainSpace. In addition to the theoretical and practical guidelines provided here, an expandable documentation has been published at http://brainspace.readthedocs.io, providing further guidance and use-case examples. BrainSpace is a simple and modular package accessible to beginners, yet expandable for advanced programmers. Core to the toolbox is a simple object-oriented model allowing for flexible computation of different i) affinity matrices, ii) dimensionality reduction techniques, iii) alignment functions, and iv) null models. As a side, we supplied precomputed gradients, a novel subparcellation of the Desikan Killani atlas, and a literature-based atlas of cortical laminar differentiation that we used in a recent study (Paquola et al., 2019; Mesulam, 1998).

As the main purpose of this report was to provide an accessible introduction of the toolboxs basic functionality, we focused on tutorial examples and several selected assessments that demonstrate more general aspects of gradient analyses First, we observed relatively consistent FC gradients were produced by multiple different dimensionality reduction techniques supplied (i.e., PCA, LE, DME), we observed relatively consistent gradients, at least when cosine similarity was chosen for affinity matrix computations. Potential interaction between input modality, affinity matrix formulation, and dimensionality reduction techniques may nevertheless occur and be worthwhile to further explore, as this might also help to understand potential differences in results. Second, while we could show a relative consistency of FC gradients across spatial scales in the case of vertexwise analyses and when parcellations with 300 nodes or more were used, we observed an interaction between the type of input data and parcellation-substrate at lower spatial scales. In fact, lower resolution structural parcellations might not capture fine-grained functional boundaries, specifically in heteromodal and paralimbic association cortices which may be less constrained by underlying structural features (Paquola et al., 2019). It will be informative for future work to clarify how input modality (e.g. FC, MPC, or diffusion MRI), the choice of parcellation, and the spatial scale impact on the spatial features identified by gradient analyses.

There are two broad ways through which the gradient method, as well as the Brainspace toolbox, may improve our understanding of neural organization and its associated functions. One avenue is the identification of similarities and differences in gradients derived from different brain measures. To address associations between cortical microstructure and macroscale function, a previous study (Paquola et al., 2019) demonstrated that gradients derived from 3D histology and myelin-sensitive MRI measures show both similarities and differences from those derived from resting state-state functional connectivity analysis (Margulies et al., 2016). This raises the possibility that the gradient method may help quantify common and distinct influences on functional and structural macroscale organization and shed light on the neural basis of more flexible (i.e., less structurally constrained) aspects of human cognition (Paquola et al., 2019).Another way that gradients can inform our understanding of how functions emerge from the cortex is through the analysis of how macroscale patterns of organization change in disease. One recent study (Hong et al., 2019), for example, demonstrated differences in the principal functional gradient, identified by Margulies et al. (2016), between individuals with autism spectrum disorder and typically developing controls. In this way, manifold-derived gradient analyses hold the possibility to characterize how macroscale functional organization may become dysfunctional in atypical neurodevelopment. To achieve both of these goals, it is important to consider gradient alignment of different gradients, and this can be achieved by a number of methods we supplied in the toolbox, such as Procrustes rotation. This allows researchers to both compare gradients from different modalities and to homogenize measures across subjects, while minimizing the changes to individual manifolds. Our across-subjects evaluations highlighted an increase in correspondence between individual subjects and the template manifold when Procrustes alignment was used compared to unaligned approaches, mainly driven by reasonably trivial changes in gradient, such as a change in the sign of specific gradients in a subgroup of subjects. As an alternative to the Procrustes alignment, it is also possible to align gradients by applying joint manifold alignments, often referred to as joint-embeddings (Xu et al., 2019).The rotations provided by embedding alignments can augment both cross-subject (Nenning et al., 2017) as well as cross-species analyses (Xu et al., 2019). Of note, this joint embedding technique generates a new manifold from the mapping between different gradients, which may result in new solutions that do not fully correspond to the initial gradients in important, and in a potentially important way.

When assessing the significance of correlations between gradients and other features of brain organization, there is an increasing awareness to ideally also evaluate correlations against null models with a similar spatial autocorrelations as the the original features. In our toolbox, we present two different approaches for building such null models, including a spin permutation test that is an adaptation of a previously released approach (Alexander-Bloch et al., 2018) and Moran’s spectral randomization used in prior ecological studies (Cliff and Ord, 1973). Gradients can also be used as a coordinate system, and stratify cortical features that are not per se gradient-based. Examples include surface-based geodesic distance measures from sensory-motor regions to other regions of cortex (Margulies et al., 2016), task-based functional activation patterns and meta-analytical data (Murphy et al., 2018, 2019; Margulies et al., 2016), as well as MRI-based cortical thickness and PET-derived amyloid beta uptake measures (Lowe et al., 2019). As such, using manifolds as a new coordinate system (Huntenburg et al., 2018) may complement widely used parcellation approaches (Schaefer et al., 2017; Yeo et al., 2011; Glasser et al., 2016) and be can be useful for dimensionality reduction of findings and for the interpretation and communication of results.

## Acknowledgements

Mr. Vos de Wael was funded by the Savoy Foundation for Epilepsy Research. Dr. Oualid Benkarim was funded by a Healthy Brains for Healthy Lives (HBHL) postdoctoral fellowship. Dr. Jessica Royer was supported by a Canadian Open Neuroscience Platform (CONP) fellowship. Dr Smallwood was supported by the European Research Council (WANDERINGMINDS – ERC646927). Dr. Paquola was funded through a postdoctoral fellowship of the Transforming Autism Care Consortium (TACC) and the Fonds de la recherche du Québec - Santé (FRQ-S). Dr. Boris Bernhardt acknowledges research support from the National Science and Engineering Research Council of Canada (NSERC, Discovery-1304413), the Canadian Institutes of Health Research (CIHR, FDN-154298), the Azrieli Center for Autism Research of the Montreal Neurological Institute (ACAR), SickKids Foundation (NI17-039), and received salary support from FRQS (Chercheur Boursier Junior 1).

## Appendix A. Python code samples

In this appendix we show the Python code analogous to the Matlab code samples provided in the main body of the paper.

Sample Code 5: Python version for Matlab code sample 1. A minimal example for plotting the first gradient of an input data matrix on the cortical surface.

~~~
from brainspace.gradient import GradientMaps
from brainspace.plotting import plot_hemispheres
from brainspace.mesh import mesh_io as mio
*# Create a GradientMaps object*
G = GradientMaps(kernel=‘cosine’, approach=‘dm’)
*# Apply GradientMaps to the data*
G.fit(data_matrix)
*# Load surfaces*
left_surface = mio.load_surface(‘left_surface_file.obj’)
right_surface = mio.load_surface(‘right_surface_file.obj’)
*# Plot first gradient on the cortical surface*
plot_hemispheres(left_surface, right_surface, G.gradients_[:,0])
~~~

Sample Code 6: Python version for Matlab code sample 2. A minimal example for creating and plotting gradients from different modalities, with different alignments.

~~~
from brainspace.gradient import GradientMaps
from brainspace.plotting import plot_hemispheres
from brainspace.mesh import mesh_io as mio
*# Create two GradientMaps objects with different alignments*
Gp = GradientMaps(kernel=‘cosine’, approach=‘dm’,
alignment=‘procrustes’)
Gj = GradientMaps(kernel=‘cosine’, approach=‘dm’,
alignment=‘joint’)
*# Apply GradientMaps to the data*
Gp.fit([mpc, fc])
Gj.fit([mpc, fc])
*# Load surfaces*
left_surface = mio.load_surface(‘left_surface_file.obj’)
right_surface = mio.load_surface(‘right_surface_file.obj’)
*# Plot first MPC gradient of Procrustes alignment*
plot_hemispheres(left_surface, right_surface, Gp.aligned_[0][:,0])
*# Plot first FC gradient of joint embedding*
plot_hemispheres(left_surface, right_surface, Gj.aligned_[1][:,0])
~~~

Sample Code 7: Python version for Matlab code sample 3. A minimal example for aligning the gradients of two individuals to a template gradient.

~~~
from brainspace.gradient import GradientMaps
from brainspace.plotting import plot_hemispheres
from brainspace.mesh import mesh_io as mio
*# Create a GradientMaps object for the template*
Gt = GradientMaps(kernel=‘cosine’, approach=‘dm’)
*# Apply to template data*
Gt.fit(template_data)
*# Create a GradientMaps object for the individuals*
Gs = GradientMaps(approach=‘dm’, kernel=‘cosine’,
alignment=‘procrustes’)
*# Compute gradients for all subjects and align to template*
Gs.fit([subject1_data, subject2_data], reference=Gt.gradients_)
*# Load surfaces*
left_surface = mio.load_surface(‘left_surface_file.obj’)
right_surface = mio.load_surface(‘right_surface_file.obj’)
*# Plot the first aligned gradient for subject 2*
plot_hemispheres(left_surface, right_surface, Gs.aligned_[1][:, 0])
~~~

Sample Code 8: Python version for Matlab code sample 4. A minimal Python example for building null models based on spin tests.

~~~
from brainspace.null_models import SpinPermutations
from brainspace.mesh import mesh_io as mio
*# Load spheres*
left_sphere = mio.load_surface(‘left_sphere_file.obj’)
right_sphere = mio.load_surface(‘right_sphere_file.obj’)
*# Generate 1000 spin permutations*
n_perm = 1000
sp = SpinPermutations(n_rep=n_perm)
sp.fit(left_sphere, points_rh=right_sphere)
features_spin = sp.randomize(left_features, x_rh=right_features)
~~~

## Footnotes

2 https://github.com/scikit-learn/scikit-learn

3 https://github.com/satra/mapalign

4 https://xaviergp.github.io/littlebrain/

5 https://github.com/koenhaak/congrads

